# Immortalized bovine satellite cells for cultured meat applications

**DOI:** 10.1101/2022.12.02.518927

**Authors:** Andrew J. Stout, Miles J. Arnett, Kristin M. Chai, Tina Guo, Lishu Liao, Addison B. Mirliani, Miriam L. Rittenberg, Michelle Shub, Eugene C. White, John S. K. Yuen, Xiaoli Zhang, David L. Kaplan

## Abstract

For cultured meat to succeed at scale, muscle cells from food-relevant species must be expanded *in vitro* in a rapid and reliable manner to produce millions of metric tons of biomass annually. Toward this goal, genetically immortalized cells offer substantial benefits over primary cells, including rapid growth, escape from cellular senescence, and consistent starting cell populations for production. Here, we develop genetically immortalized bovine satellite cells (iBSCs) via constitutive expression of bovine Telomerase reverse transcriptase (TERT) and Cyclin-dependent kinase 4 (CDK4). These cells achieve over 120 doublings at the time of publication and maintain their capacity for myogenic differentiation. They therefore offer a valuable tool to the field, enabling further research and development to advance cultured meat.

## Introduction

The production of cultured meat, poultry, and seafood products through cell culture and tissue engineering will depend on unprecedented levels of animal cell expansion *in vitro* in order to compete with conventional meat products. For instance, global beef consumption is expected to grow to 75 million metric tons by 2030, part of an upward trend which has continued since at least 1961^1,2^. In order to produce sufficient biomass *in vitro* to meet this demand, muscle and fat cells will need to have high proliferative capacity, both in terms of doubling rate and in terms of the total number of achievable doublings. For the latter, primary stem cells obtained directly from animal tissue are typically limited to ∼50 doublings, a threshold known as the Hayflick limit^3^. During these doublings, a cell’s telomeres (protective regions at the ends of chromosomes) are progressively shortened due to incomplete replication by the cell’s polymerases. Upon full depletion of these telomeres (the Hayflick limit), further chromosome shortening results in DNA damage which induces cell cycle arrest, morphological changes, and a range of other cellular changes known as senescence^4^.

While it is theoretically possible for a relatively small number of starting primary cells to produce a large amount of biomass after fifty doublings^5^, there are several reasons why immortalized cells, engineered or adapted to bypass senescence, offer benefits to cultured meat research and production. For instance, the need to obtain primary cell populations requires research groups to source tissues, establish or validate isolation protocols, and characterize cell populations before they can begin meaningful research with primary cells. Alternatively, some groups rely on model cell lines, such as the immortalized C2C12 mouse muscle cell line, which are useful in studies but ultimately offer limited direct analogy to meat-relevant species for consumers. For commercial production, the use of primary cells would require repeat tissue biopsies from animals which could affect process reproducibility due to variations in animal biology or isolated cell populations^6^. Additionally, further engineering or adapting primary cells to optimize cultured meat products (e.g., through enhanced nutrition or tolerance for waste metabolites) is impractical at production scale, due to the need to re-engineer each new biopsied cell population^7,8^. Thus, immortalized cells are required for further cell engineering and optimization, which would have substantial value for cultured meat production processes. Finally, apart from senescence-induced cell cycle arrest, the myogenic capacity of primary muscle satellite cells decreases significantly over prolonged culture, and is substantially reduced well before the Hayflick limit of 50 doublings^6,9^. This is in contrast with immortalized muscle cells, which retain their differentiation capacity indefinitely^10^.

To date, numerous strategies have been explored to immortalize myogenic precursor cells. These include spontaneous immortalization whereby random mutations accumulated during serial culture resulting in outgrowth of an immortal population^11^, infection with simian virus 40 (SV40), ectopic expression of the SV40 Large T antigen^12,13^, or ectopic expression of telomerase reverse transcriptase (TERT) and Cyclin dependent kinase 4 (CDK4)^10,14^. In the latter case, TERT acts by elongating telomeres to counteract telomere shortening, and CDK4 promotes progression through the G1/S phase transition, thereby providing an “engine” for cell division. This approach has been demonstrated for human satellite cells for the study of muscular disease^10,14–16^, as well as fibroblasts from various species (e.g., marmoset, bovine, porcine, rabbit, and others) for the purpose of general research and species preservation^17–20^. However, to our knowledge, it has not yet been applied to satellite cells from food-relevant species.

For cultured meat applications, the TERT/CDK4 approach offers the advantages of being consistently reliable (unlike spontaneous immortalization approaches) and non-oncogenic (unlike SV40), which could offer benefits from consumer and regulatory standpoints. While this approach has been demonstrated for myogenic cells from humans and mice, it has not yet been applied to cells relevant to cultured beef. In the present work, primary bovine satellite cells (BSCs) are engineered via transposon integration of TERT and CDK4 to establish immortalized bovine myogenic cells (iBSCs). These iBSCs offer robust proliferative capacity (>120 doublings; 15 hours per doubling) while maintaining the capacity to fuse into multinucleated myotubes, albeit to a reduced degree compared with primary BSCs. Together, this work demonstrates the efficacy of TERT/CDK4 expression in immortalizing bovine muscle progenitors and offers a valuable tool for cultured meat research and development.

## Results

### Starting materials: Primary bovine satellite cells & the TERT/CDK4 transposon system

Primary BSCs used in this study were characterized for their phenotype and myogenicity. First, proliferative cells were analyzed for expression of the early-stage satellite cell identity marker Paired-box 7 (Pax7) and the early myogenic activation marker Myoblast Determination Protein 1 (MyoD). Immunostaining revealed that the BSCs showed ubiquitous Pax7 expression, and heterogenous but consistent MyoD expression (Figure 1a). Upon reaching confluence, cells were differentiated by incubating cells for six days without media changes, and differentiated cells were analyzed for expression of the early differentiation marker Myogenin and the terminal fusion marker Myosin heavy chain (MHC). The results showed that BSCs differentiated into multinucleated myotubes which were positive for both Myogenin and MHC (Figure 1b). Together, these results validated the identity and myogenicity of primary BSC populations.

**Figure 1:**
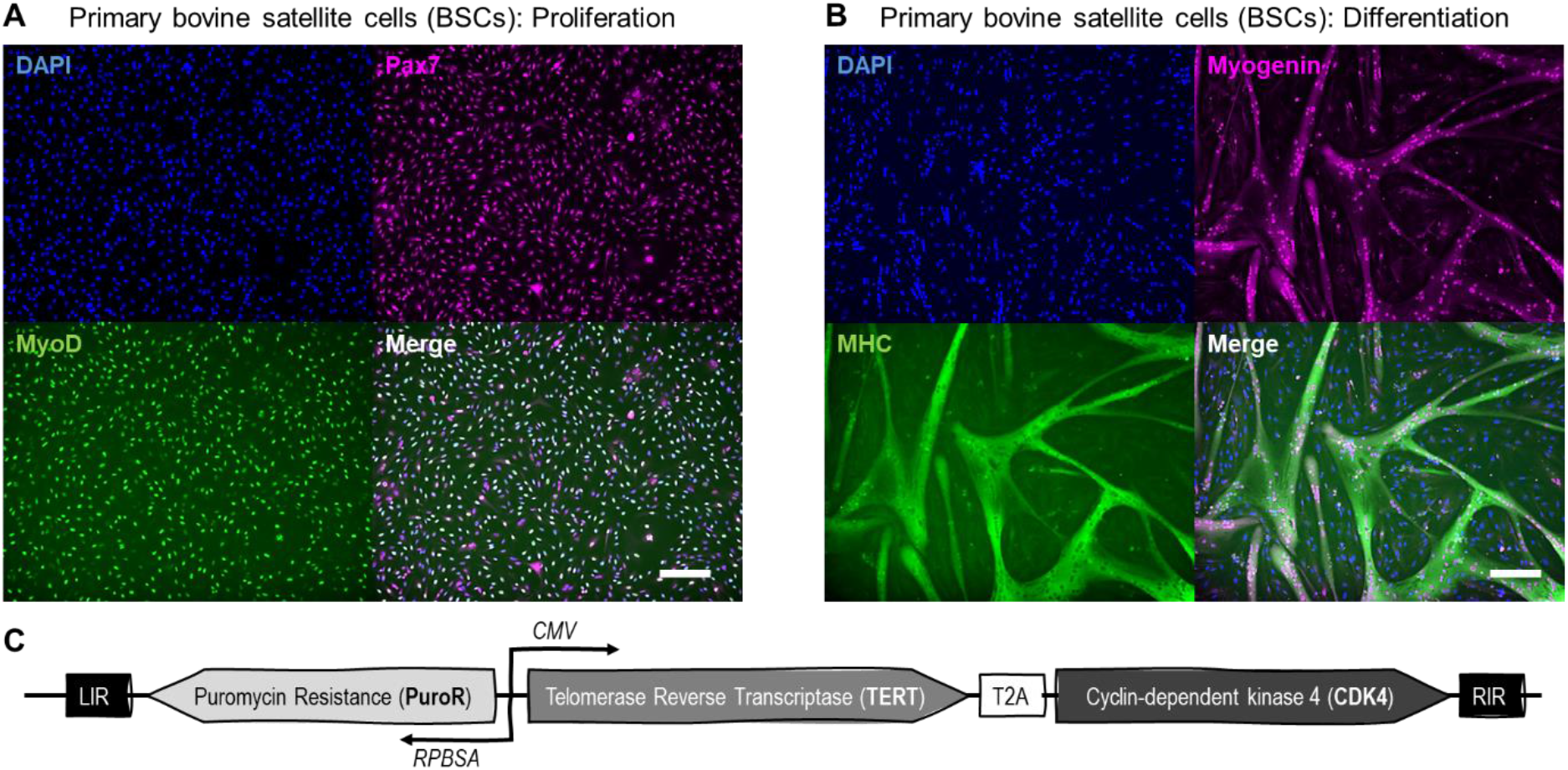
Starting cells and genetic constructs. (A) Primary bovine satellite cells (BSCs) were cultured to 70% confluency and stained for two markers of cell identity: Paired-box 7 (Pax7; magenta), the early satellite cell marker, and Myoblast determination protein 1 (MyoD; green), a marker of myogenic commitment. A nuclear counterstain (DAPI; blue) was also used. The results showed ubiquitous Pax7 expression and MyoD expression in all nuclei, with noticeably heterogeneous expression of MyoD. Scale bar is 200 μm. (B) Upon differentiation for two days in 2% FBS-containing medium, BSCs fused into multinucleated myotubes which stained positive for the early differentiation marker myogenin (magenta) and the terminal fusion marker Myosin heavy chain (MHC; green), indicating robust myogenic capacity of the primary cells. Scale bar is 200 μm. (C) A schematic of the *Sleeping Beauty* transposon system that was used to engineer BSCs for constitutive expression of telomerase reverse transcriptase (TERT) and cyclin-dependent kinase 4 (CDK4), as well as a puromycin resistance element (PuroR).

Once the starting BSCs had been characterized, the cells were engineered through *Sleeping Beauty* transposon-mediated insertion of bovine TERT and bovine CDK4 genes under the cytomegalovirus (CMV) promoter, as well as the gene for puromycin resistance under a synthetic RPBSA promoter (Figure 1c, Figure S1)^21^. A T2A linker was used to enable near-stoichiometric bi-cistronic expression of TERT and CDK4^22^.

### Engineered cells demonstrate an immortalized muscle precursor phenotype

Following puromycin selection and cell recovery (ten passages after transfection), long-term growth was assessed for engineered cells compared with primary BSCs. The results showed that BSCs + TERT/CDK4 (iBSCs) continued to expand consistently for more than 120 doublings, while the unmodified primary BSCs began to slow after approximately 30 doublings, and effectively stopped dividing by the 45th doubling (Figure 2). This indicates that iBSCs evaded cellular senescence in contrast to the primary BSCs, crossing the 100-doubling threshold that is often used to define successful immortalization^23–26^.

**Figure 2:**
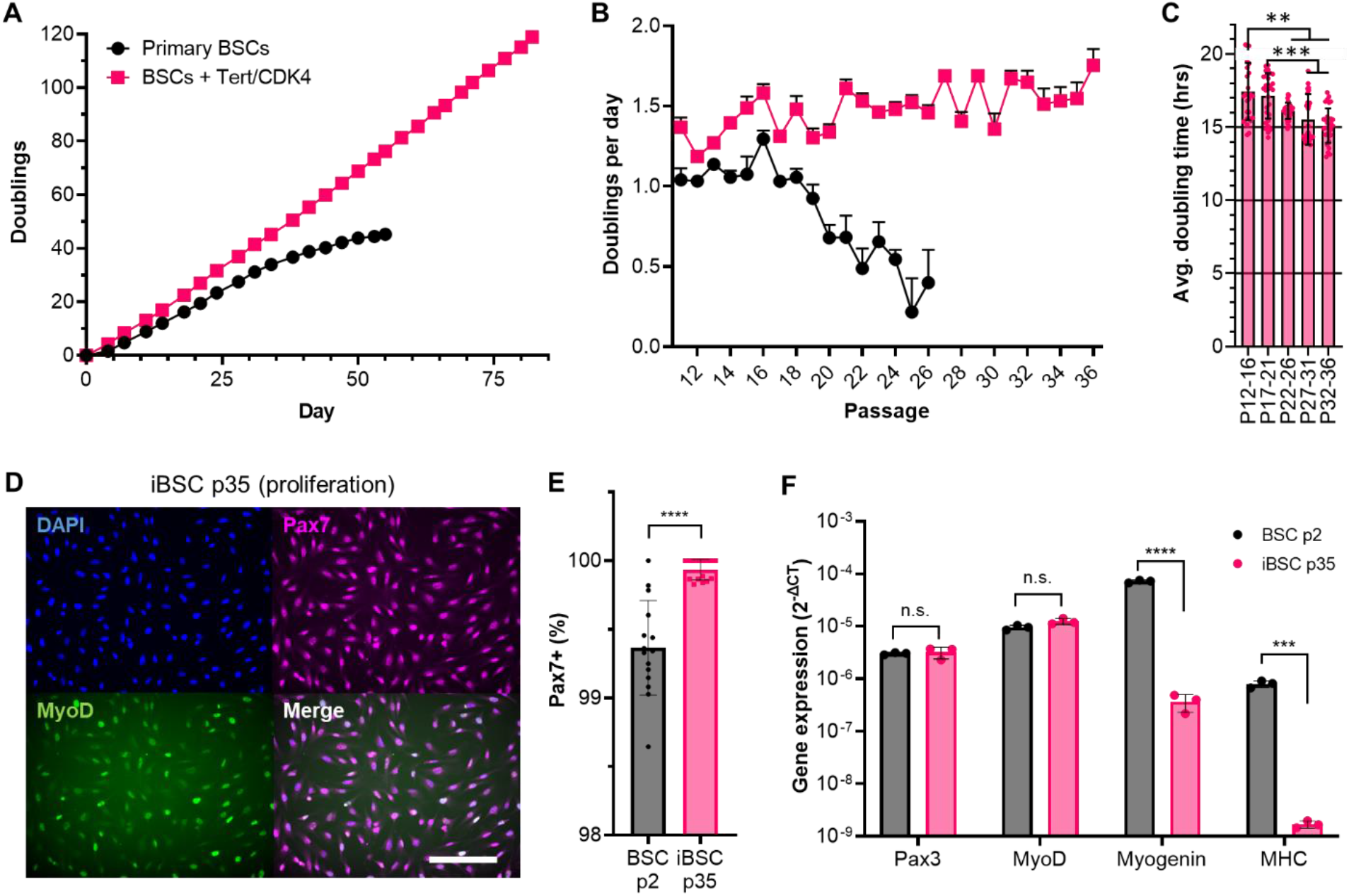
Long-term growth of engineered BSCs. (A) Cumulative population doublings of unmodified primary BSCs and BSCs engineered with TERT and CDK4 (iBSCs). The results showed that iBSCs maintained consistent growth over >120 doublings, while primary BSCs lost growth capacity around 45 doublings. (B) Doubling rate for primary and immortalized cells as indicated by doublings per day. Results showed that iBSCs maintain high doubling rate, while primary BSCs lose growth rate substantially after passage 16. (C) Converting doublings per day to doubling time averaged over five-passage increments reveals signficantly accelerated growth rate for iBSCs from 17.4 hours at the start of the experiment to 15.1 hours by passage 36. n = 30 distinct samples (6 samples for each passage, averaged over five passages); statistical significance was calculated by one-way ANOVA with comparisons between all samples, and is indicated by asterisks, in which p <0.01 (**), p < 0.001 (***). (D) Immunostaining of subconfluent iBSCs for Pax7 and MyoD reveals ubiquitous Pax7 expression (magenta) and heterogeneous MyoD expression (green) for all cells (nuclie; blue). Scale bars at 200 μm. (E) Quantitative image analysis of Pax7 staining revealed a significant increase in the percent of Pax7+ cells for iBSCs compared with primary cells, though, which was >99% for both populations. (F) Gene expression analysis revealed no significant difference between primary and immortalized cells for Pax3 and MyoD expression, but a significant decrease in myogenin and MHC expression for iBSCs. As these two genes characterize a differentiated phenotype, these results suggest a more undifferentiated phenotype for immortalized cells. n = 3 distinct samples; statistical significance was calculated via paired t-tests for each gene, and is indicated by asterisks, in which p <0.05 (*), p <0.01 (**), p < 0.001 (***) and p < 0.0001 (****).

Converting growth data into doubling rate further illustrates the robust growth of the iBSCs, which maintain a consistent high doubling rate throughout the experiment (Figure 2b). This was in contrast with primary BSCs, which slowed significantly after the 16^th^ passage. Interestingly, the growth rate of the iBSCs increased over the course of the experiment as the average doubling time of iBSCs fell from 17.4 hours during early passages (P12-P16) to 15.1 hours during later passages (P32-P36) (Figure 2c). This acceleration could be due to the inherent selection for faster-growing subpopulations, though whether further acceleration is possible over longer culture periods remains to be seen. The final growth rate of 15 hours per doubling is noteworthy in the context of large-scale cell culture for cultured meat, as this is faster than the ∼17 hours for BSCs *in vivo*, and on-par with growth rates for traditional bioprocess cell lines like Chinese hamster ovary (CHO) cells^27,28^.

Following successful immortalization, phenotypic analysis of proliferating cells demonstrated that iBSCs maintained their stem cell identity over long-term passaging (Figure 2d-f). Specifically, immunostaining of iBSCs after 35 passages revealed ubiquitous Pax7 expression and heterogeneous MyoD expression similar to the primary BSCs in Figure 1. Quantitative image analysis of Pax7 staining further supported this conclusion, showing that >99% of both primary BSCs (P2) and immortalized iBSCs (P35) were positive for the marker. Next, quantitative PCR (qPCR) analysis of gene expression was performed to further understand the proliferative iBSC phenotype compared with primary BSCs. The results showed that, for early satellite cell markers Pax3 and MyoD, no significant difference existed between the primary and immortalized cells. For differentiation markers myogenin and myosin heavy chain (MHC), however, the results showed a significant reduction in gene expression for the immortalized iBSCs compared with primary BSCs. This suggests that iBSCs might be able to maintain a more undifferentiated phenotype than primary cells, which could be associated with their enhanced proliferative capacity. Together, these results indicated that the iBSCs maintained the BSC phenotype even after extensive passaging, suggesting that cell engineering and immortalization did not alter the stem cell identity of iBSCs.

### iBSCs maintain their myogenic capacity

Once the iBSC line was established and its stem cell phenotype was confirmed, the cells were assessed for their ability to differentiate into multinucleated myotubes. Specifically, iBSCs and primary BSCs were cultured to confluency, differentiated as before (by changing to a 2% FBS differentiation medium), and assessed for the formation of multinucleated myotubes. The iBSCs generated myotubes at a reduced rate when compared to the primary BSCs, thus the BSCs were differentiated for two days before further analysis, while the iBSCs were differentiated for both two and three days for comparisons. Immunostaining revealed that the iBSCs retained their ability to differentiate into multi-nucleated myotubes (Figure 3a, Figure S2), and that differentiated cells expressed both myogenin and MHC. These results indicated that the iBSCs maintained their myogenic phenotype.

**Figure 3:**
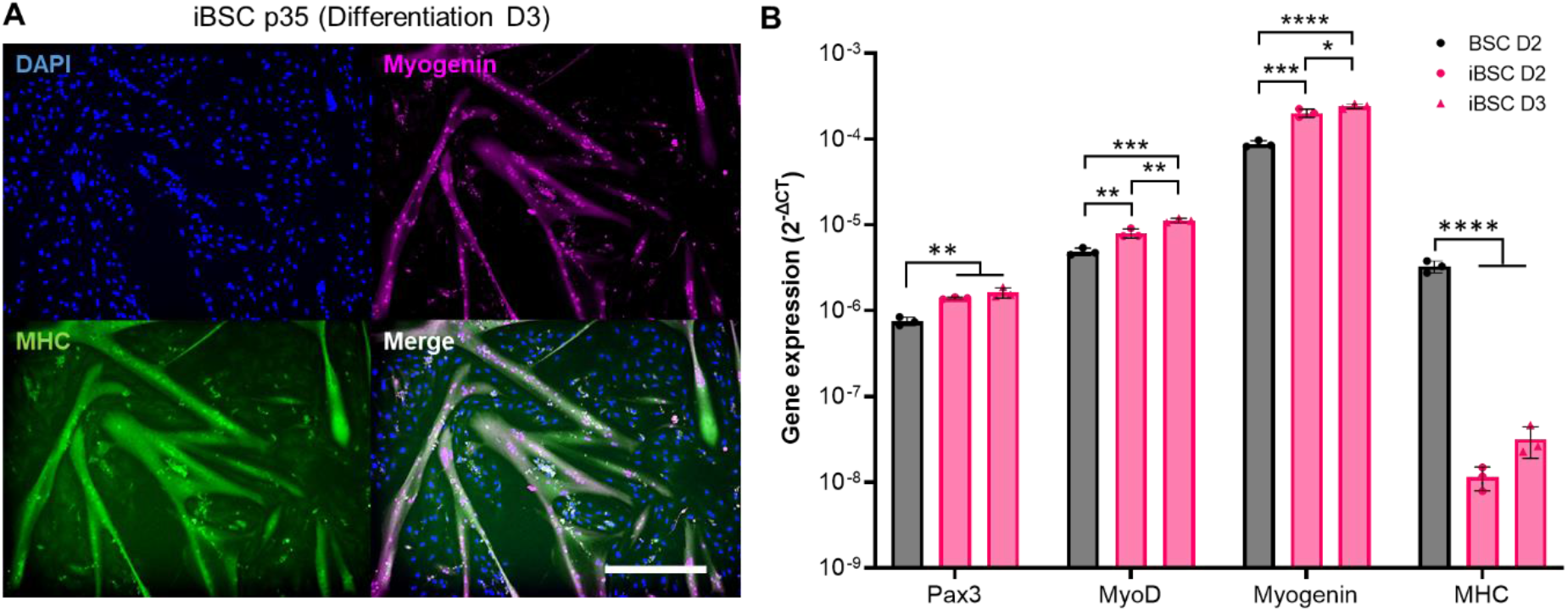
Differentiation of iBSCs. (A) Immunostaining of iBSCs differentiated for three days in 2% FBS medium and stained for nuclie (DAPI; blue) myogenin (magenta) and MHC (green). The results showed the robust formation of MHC-positive multinucleated myotubes containing nuclei that were positive for myogenin. Staining and differentiative phenotype is similar to that seen for primary BSCs in Figure 1, which confirms that iBSCs maintain the myogenic phenotype of these cells. (B) Gene expression analysis of differentiated primary BSCs on day two of differentiation (D2) and of iBSCs on days two (D2) and three (D3) of differentiation. The results showed a significant increase in early-stage myogenic markers (Pax3, MyoD, and Myogenin) for iBSCs, but a significant decrease in the late-stage maker MHC. A significant increase in MyoD and Myogenin (and a non-significant trend upwards in MHC) was seen with increased differentiation time of iBSCs. n = 3 distinct samples; statistical significance was calculated by one-way ANOVA for each gene, with comparisons between all samples and is indicated by asterisks, in which p <0.05 (*), p <0.01 (**), p < 0.001 (***) and p < 0.0001 (****).

Comparing gene expression between the differentiated primary and immortalized cells showed that iBSCs offered significantly higher expression of pax3, MyoD, and myogenin, but significantly reduced expression of MHC (Figure 3). These results were in line with the observed speed of differentiation for the iBSCs compared with the primary BSCs and suggested that the more undifferentiated iBSC phenotype observed in Figure 2 might also impact cellular differentiation. Specifically, iBSCs demonstrated a reduced degree of terminal differentiation. While no significant difference was seen for MHC expression between two and three-day iBSC differentiation, a significant increase was observed for MyoD and myogenin. These data, along with the MHC trends, suggested that increased culture time could enhance the degree of differentiation in iBSCs. Together, these results suggested that iBSCs maintained their capacity for myogenic differentiation but at a reduced level than primary BSCs under similar conditions and time periods.

### iBSCs show enhanced TERT/CDK4 expression and heterogenous chromosomal integration

Following validation of the iBSCs as an immortalized myogenic bovine cell line, the nature of transgene expression and integration was assessed. First, qPCR was performed to compare TERT and CDK4 expression in the unmodified and engineered cells (Figure 4). The results showed that, while primary BSCs had substantially higher native CDK4 expression than TERT expression, both genes were significantly upregulated in the iBSCs, indicating successful transgene insertion and ectopic expression. Next, karyotyping was performed to validate that immortalized cells showed a normal female bovine karyotype (Figure 4b, Figure S3). Finally, possible integration sites were assessed. The results showed potential insertion sites of TERT and CDK4 at chromosomes 18, 21 and 29, with the integrated transposons flanked by the TA boundaries characteristic of the *Sleeping Beauty* system (Figure 4b). Together, these results show that the TERT/CDK4 immortalization construct was integrated at multiple points in the chromosomes, and that this integration resulted in successful overexpression of both genes, leading to the observed cellular immortalization.

**Figure 4:**
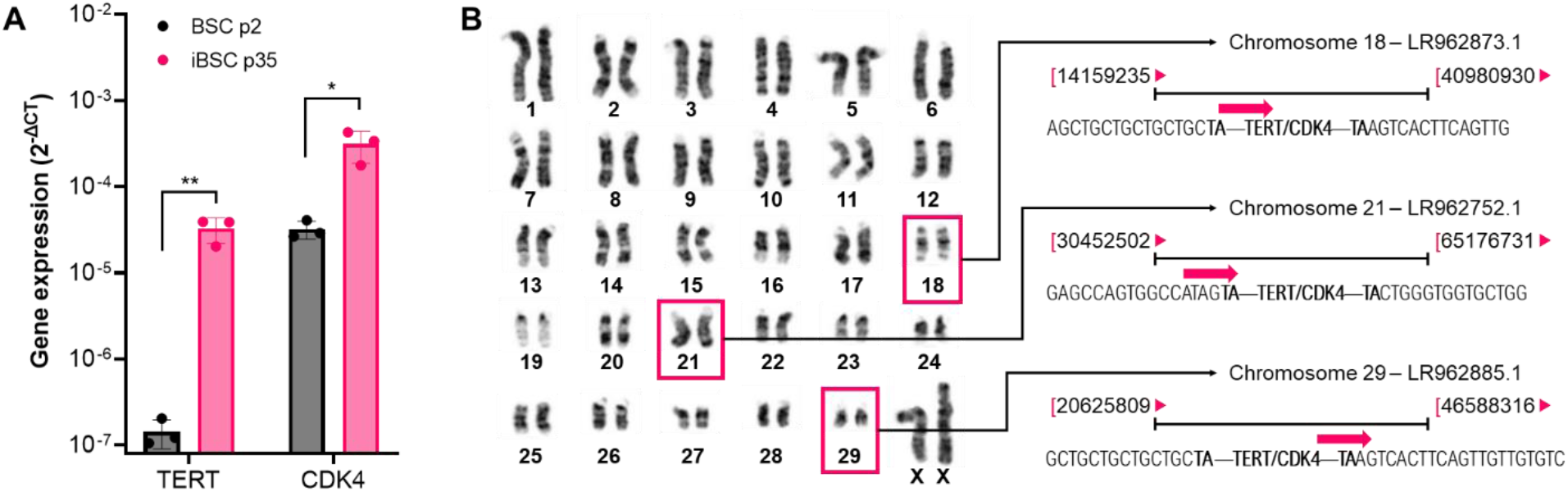
Transgene characterization. (A) Gene expression analysis revealed a significant increase in expression of TERT and CDK4 cells in engineered cells compared with unmodified primary BSCs. (B) Karyotype analysis revealed a normal female karyotype, and integration analysis revealed three potential insertion locations for transposed TERT/CDK4 constructs, including in chromosomes 18, 21, and 29 of the cattle genomes. In all cases, TERT/CDK4 constructs were flanked by characteristic TA dinucleotides.

## Discussion

The success of cultured meat will depend on cell populations that can expand quickly, reliably, and in low-cost environments (e.g., low-cost culture media). The ability of these cells to be engineered or adapted to low-cost bioprocesses would further enable compounding benefits, allowing for the development of optimized and consistent cell lines for cultured meat production. Immortalized progenitor cells offer a valuable avenue for generating such cell lines, though no lines of muscle precursor cells from relevant species (e.g., cattle) have previously been generated. In this work, well-documented myoblast immortalization techniques were applied to bovine satellite cells in order to generate immortalized bovine myogenic cells (iBSCs). These cells demonstrate robust and rapid proliferation while retaining their myoblast phenotype and the capacity for myogenic differentiation. In doing so, these cells offer a valuable research tool to the field of cultured meat by lowering the barrier to entry for researchers while providing insight which can inform future cell engineering, bioprocess design, and techno-economic or life-cycle analyses.

An interesting outcome of this work was the speed of cell doubling for iBSCs, which reached a frequency of once every 15 hours by the time cells had surpassed 100 doublings. This is in comparison with ∼17 hours for BSCs *in vivo* or ∼24 hours for primary BSCs *in vitro* averaged over five passages (a rate which rapidly declines throughout the course of BSC growth)^9,27^. The iBSCs show a growth rate similar to biopharmaceutical workhorse cells such as the Chinese hamster ovary (CHO) cell line (∼14-17 hours per doubling), immortalized C2C12 mouse muscle cells (∼14 hours per doubling), and immortalized L6 rat myoblasts (∼15 hours per doubling)^28–30^.

From a consumer acceptance perspective, two factors must be considered when looking at iBSCs for cultured meat production. The first is that these cells are genetically modified (GM), involving transgene insertion of “foreign” genetic material. Genetic engineering of foodstuffs still raises substantial concern among consumers due to factors such as food neophobia, unfamiliarity with the technology, or perceived unnaturalness^31–33^. This aversion would need to be overcome for the benefits of immortalized cell cultures to be realized in cultured meat technologies. However, evidence suggests that increased information can reduce consumer preference towards non-GM foods, suggesting that such changes in consumer attitudes are possible^34^. More, it has been shown that the source of GM technologies can substantially impact consumer perceptions, and that consumers tend to look more favorably on GM technologies that were developed in non-profit or academic settings; as such, the iBSCs developed in this work may have some advantages from a consumer acceptance standpoint^31^.

The second consumer consideration relates to the fact that, *in vivo*, senescence escape and overexpression of TERT and CDK4 can be associated with various cancers^35,36^. There is to-date no evidence for DNA transfer from gene edited foods to humans^37^. Similarly, there is no compelling possible mechanism for iBSCs to survive cooking and digestion and pass through the intestinal wall to implant as a tumorigenic cell population in humans. As such, there is no apparent risk of immortalized cell consumption being cancerous. Still, the association could cause concern with consumers, and so further evidence of safety should be generated.

From a government perspective, the regulation and labeling of GM cells in cultured meat remains to be determined and will likely depend on policies in specific jurisdictions. For instance, while the United States has a relatively favorable regulatory landscape towards GM crops, and has to-date approved two genetically modified animals for human consumption^38,39^, the European Union poses much harsher regulatory controls on GM technologies^40,41^. Here, an important distinction is materializing between “genetically modified” organisms (which involve transgenic insertions of foreign DNA), and “gene edited” organisms, in which only a small mutation, deletion, or cisgenic insertions of native genes are imparted^41,42^. This distinction would favor targeted techniques over transgenic insertions, and so targeted immortalization strategies (such as CRISPR/Cas9 genome engineering) should be explored alongside the TERT/CDK4 expression system used in this study. Additionally from a regulatory perspective, the random insertion of the transposon system used herein, and the potential for mutation accumulation or genetic drift in immortalized cell lines, may offer challenges in terms of characterizing and monitoring cells^43,44^. Importantly, the United States Food and Drug Administration (FDA) recently completed its first pre-market consultation on cultured meat, including chicken produced with cells expressing cisgenic TERT, and provided a “no further questions” letter, indicating the FDA’s initial approval of such methods for cultured meat^45^. This provides a strong indication that genetically immortalized cells will be approved for consumption, at least in the United States.

While the realities of consumer and regulatory acceptance of genetically immortalized cells for cultured meat remains to be seen across populations and jurisdictions, the potential utility of these cells in research and development is noteworthy. Immortalized BSCs can lower barriers to entry for research groups, accelerate research projects, and increase the relevance of research findings. This is the case both for cultured meat, and for animal and veterinary sciences.

Ultimately, this work offers support for the use of TERT/CDK4 immortalization strategies in cultured meat-relevant muscle cells and provides a useful tool to the field.

## Materials and Methods

### Primary bovine satellite cell isolations

Primary BSCs were isolated following methods previously reported and in accordance with approved protocols (IACUC protocol #G2018-36)^9^. Briefly, a small (∼0.5 g) muscle excision was taken from the semitendinosus of a 5-week-old Simmental calf at the Tufts Cummings School of Veterinary Medicine. The tissue was transported on ice in DMEM+Glutamax (ThermoFisher #10566024, Waltham, MA, USA) supplemented with 1% antibiotic/antimycotic (ThermoFisher #1540062).

Once in a sterile biosafety cabinet, the tissue was minced to paste and incubated at 37°C in DMEM+Glutamax plus 0.2% collagenase II (Worthington Biochemical #LS004176, Lakewood, NJ, USA) for 45 minutes with trituration every 15 minutes. Once tissue could be passed through an 18-gauge needle, the digestion was halted with BSC growth medium (BSC-GM) made up of DMEM+Glutamax, 20% fetal bovine serum (FBS; ThermoFisher #26140079), 1 ng/mL human FGF-2 (PeproTech #100-18B, Rocky Hill, NJ, USA), and 1% Primocin (Invivogen #ant-pm-1, San Diego, CA, USA). Next, cells were passed sequentially through 70 μm and 40 μm cell strainers (Sigma #CLS431751-50EA; #CLS431750-50EA, Burlington, MA, USA), counted with an NC-200 automated cell counter (Chemometec, Allerod, Denmark), and plated at 100,000 cells/cm^2^ onto uncoated tissue culture flasks in a 37°C incubator with 5% CO_2_. After 24 hours of incubation, media and unadhered cells (the satellite cell fraction) were transferred to a new flask along with 0.25 ug/cm^2^ iMatrix recombinant laminin-511 (Iwai North America #N892021, San Carlos, CA, USA). Flasks were left untouched for three days before growth media was changed.

After this, cells were cultured using standard practices on tissue-culture plastic with BSC-GM (changed every 2-3 days) and 0.25 ug/cm^2^ of iMatrix laminin-511 (added each time cells were seeded onto new culture surfaces). Cells were cultured to 70% confluency, harvested with 0.25% trypsin-EDTA (ThermoFisher #25200056), and either passaged or cryopreserved in FBS with 10% Dimethyl sulfoxide (DMSO; Sigma #D2650).

### Primary BSC characterization

To assess proliferative BSCs, cells were cultured to 70% confluency, fixed for 30 minutes in 4% paraformaldehyde (ThermoFisher #AAJ61899AK), rinsed 3X with DPBS (ThermoFisher #14190250), permeabilized for 15 minutes in DPBS with 0.5% Triton-X (Sigma # T8787), rinsed 3X with PBST [DPBS plus 0.1% Tween-20 (Sigma #P1379)], and blocked for 45 minutes in blocking buffer [DPBS with 5% goat serum (ThermoFisher #16210064) and 0.05% sodium azide (Sigma #S2002)]. Primary antibodies for Pax7 (ThermoFisher #PA5-68506) and MyoD (ThermoFisher #MA5-12902) were diluted in blocking buffer 1:500 and 1:100, respectively, and incubated overnight with cells at 4°C. The next day, cells were rinsed 3X with PBST, incubated with blocking buffer for 15 minutes, and incubated for one hour at room temperature with blocking buffer containing secondary antibodies for Pax7 (ThermoFisher #A-11072; 1:500) and MyoD (ThermoFisher #A-11001), as well as a DAPI nuclear stain (Abcam #ab104139, Cambridge, UK; 1:1,000). Finally, cells were rinsed 3X with DPBS and imaged on a KEYENCE BZ-X810 fluorescent microscope (Osaka, Japan).

To assess differentiated BSCs, cells were cultured to confluency and media was changed to DMEM supplemented with 2% FBS and 1% antibiotic-antimycotic. Cells were differentiated for two days. Cells were then fixed, permeabilized, blocked, and stained with the same protocols as for proliferative cells. Primary antibodies for Myogenin (ThermoFisher #PA5-116750) and MHC (Developmental studies hybridoma bank #MF-20, Iowa City, IA, USA) were used at dilutions of 1:100 and 4 μg/mL, respectively, and DAPI and the same secondary antibodies were used as with proliferative analyses.

### Cloning and transfections

DNA for bovine TERT (Accession #NM_001046242.) and CDK4 (Accession # NM_001037594.2) were obtained as open reading frame clones with additional terminal Flag and HA tags, respectively (GenScript, Piscataway, NJ, USA). Genes were inserted into a *Sleeping Beauty* plasmid which had been previously developed by our group through the modification of the pSBbi-Pur plasmid from Eric Kowarz (Addgene #60523)^7,46^. This new plasmid, termed pSB-Imm-Puro, contained TERT and CDK4 which were separated by the sequence for a self-cleaving T2A peptide^22^, promoted by a CMV promoter, and followed by a bGH PolyA signal. A synthetic RPBSA promoter ran counter to CMV and promoted expression of a puromycin resistance gene, and an ampicillin resistance and origin of replication were present for standard plasmid maintenance. All cloning was performed via gibson assemblies, which involved fragment amplification with the Q5 high-fidelity DNA polymerase (NEB #M0494S, Ipswich, MA, USA), DpnI digestion (NEB #R0176S), PCR cleanup (NEB #T1030S), NEBuilder HiFi assembly (NEB #E2621S), and transformation into chemically competent *E. coli* (NEB #C3019H), all according to the manufacturer’s instructions. Assemblies were verified with Sanger sequencing (Genewiz, Cambridge, MA, USA), and plasmids were purified via GeneJet miniprep (ThermoFisher #K0503). The transposase plasmid pCMV(CAT)T7-SB100 was a gift from Zsuzsanna Izsvak (Addgene #34879), and was similarly purified^47^.

For transfecting BSCs, cells were thawed and seeded at 25,000 cells/cm^2^ in 60well plates with BSC-GM and iMatrix laminin-511. The next day, cells were transfected with Lipofectamine 3000 (ThermoFisher #L3000015) according to the manufacturer’s protocol adapted for multi-plasmid transfection. Briefly, 2.5 μg of pSB-Imm-Puro was combined with 0.25 μg of pCMV(CAT)T7-SB100 in 250 uL of Opti-MEM medium (ThermoFisher #31985088) with 7.5 uL of Lipofectamine 3000, and 5 uL of p3000. The DNA solution was incubated for 15 minutes at room temperature, during which time cells were rinsed 1X with DPBS and fed 2 mL of Opti-MEM. After incubation, the DNA solution was added to the cells, and cells were incubated for six hours at 37°C before 2 mL of BSC-GM was added to wells containing Opti-MEM and lipofectamine mixture. Cells were incubated at 37°C overnight, after which media was replaced with BSC-GM supplemented with 2.5 μg/mL puromycin (ThermoFisher #A1113803). For the remainder of the study, engineered cells were cultured in this puromycin-supplemented media according to the same protocols as for primary BSCs.

### Long-term growth curves

Once engineered cell growth had stabilized under puromycin selection (10 passages), long-term growth was analyzed through serial passaging. Briefly, iBSCs and primary BSCs were seeded in triplicate in 6-well plates at 20,000 cells/well (2,083 cells/cm^2^) with BCS-GM with iMatrix laminin-511 and ± puromycin. Media was changed every other day until cells reached 70-80% confluency, at which point cells were harvested, counted (with technical duplicates for cell counts) and re-seeded in new 6-well plates as described above. Every five passages, a subset of cells was frozen in FBS + 10% DMSO. Primary BSC culture was halted after cell growth drastically slowed, indicating that senescence had been reached. After 120 doublings had been achieved for engineered cells (passage 35 from the initial isolation), these were designated as immortalized bovine myogenic cells (iBSCs).

### Immunostaining and quantitative image analysis of iBSCs

Once established, iBSCs (≥passage 35) were characterized and compared with primary BSCs (passage 2) for phenotypic markers and myogenic capacity. First, proliferative cells (P35) were fixed and stained for DAPI, Pax7 and MyoD as previously described. Next, cells were cultured to confluency, and differentiated in DMEM + 2% FBS for either two or three days before being fixed and stained for DAPI, Myogenin and MHC as previously described. For quantitative Pax7 analysis, batch images were taken at five random points within triplicate wells of cells using the microscope’s random point selection tool. Images were then analyzed in ImageJ by gating regions of interest using the DAPI channel and determining the percentage of thus-gated nuclei that contained Pax7 signal after a consistent threshold was applied.

### Gene expression analysis

Relative gene expression between proliferative and differentiated BSCs and iBSCs was assessed via quantitative PCR using the TaqMan Fast Universal PCR Master Mix without AmpErase UNG (ThermoFisher #4352042). Briefly, RNA was harvested at 70% confluency or after differentiation using the RNEasy Mini kit (Qiagen #74104, Hilden, Germany) according to the manufacturer’s instructions. Next, cDNA was prepared using the iScript cDNA synthesis kit (Bio-Rad #1708890, Hercules, CA, USA) and 750 ng of RNA for each reaction. Finally, qPCR was performed using 2 uL of cDNA and primers for 18S (ThermoFisher #Hs03003631), Pax3 (ThermoFisher #Bt04303789), MyoD1 (ThermoFisher #Bt03244740), Myogenin (ThermoFisher #Bt03258929), Myosin Heavy Chain (ThermoFisher #Bt03273061), TERT (ThermoFisher #Bt03239211), and CDK4 (ThermoFisher #Bt03231354). Reactions were performed on a CFX96 Real Time System thermocycler (Bio-Rad, Hercules, CA, USA), and results were analyzed as 2^-Δct^ normalized to expression of the 18S housekeeping gene.

### Karyotyping and chromosomal integration analysis

Karyotyping was performed by Creative Bioarray (Sherley, NY, USA) on iBSCs at passage 28. Twenty G-banded metaphase spreads were analyzed using a GTG Banding technique. To analyze chromosomal integration, genomic DNA of the iBSC was extracted via GeneJet Genomic DNA purification kit (ThermoFisher #K0721) according to the manufacturer’s instructions. To detect the chromosome location of integration, primers (pSB-3’-F: GCTCCTAACTGACCTTAAGACAGGG; pSB-5’-R: GCTCTGACCCACTGGAATTGTGA) were designed based on the IR-DR Sequence of the sleeping beauty transposon vector^48^, and PCRs were performed via Phusion® High-Fidelity PCR Master Mix with HF Buffer (NEB #M0531S). After PCR clean-up (Promega #A9281, Madison, WI, USA), A-tailing of the PCR products with Taq Polymerase was performed as: 1) reaction set-up with PCR-amplified DNA, 5μL of 10x ThermoPol® Buffer (NEB #B9004), 10 μL of 1mM dATP, 0.2 μL of Taq DNA Polymerase (NEB #M0267), and ddH2O to a final volume of 50 μL; 2) Incubation at 72°C for 20 min. The A-tailed PCR products were then cloned into the pGEM®-T Easy Vector Systems (Promega #A1360) followed by transformation. After mini-prepping (ThermoFisher #IB47101), plasmid clones were digested by EcoRI-HF (NEB #R3101S) to confirm the insertion, and then sent for sequencing. The sequence results were investigated using NCBI Blast to identify candidate chromosome locations of the transposons.

### Statistical analysis

Statistical analysis was performed with GraphPad Prism 9.0 software (San Diego, CA, USA). Doubling time and differentiated gene expression were analyzed via one-way ANOVA. Multiple comparisons were performed with Tukey’s HSD post-hoc test with comparisons between all samples. Pax7+ percentage and proliferative gene expression were analyzed via unpaired t-tests. P values <0.05 were treated as significant. All error bars represent ± standard deviation.

## Supporting information

Suppelementary Materials

## Author contributions

**AJS**: Conceptualization, Methodology, Investigation, Formal Analysis, Visualization, Writing-original draft preparation, Writing-reviewing and editing. **MJA**: Investigation, Formal Analysis, Writing-original draft preparation. **KC**: Investigation. **TG**: Investigation, Formal Analysis. **ABM**: Investigation, Formal Analysis. **MLR**: Investigation. **MS**: Investigation. ECW: Methodology, Resources. **JSKY, Jr**.: Conceptualization, Methodology. **XZ**: Investigation, Methodology, Formal Analysis, Writing-original draft preparation. **DLK**: Conceptualization, Resources, Writing-reviewing and editing, Supervision, Funding acquisition.

## Acknowledgements

We thank the USDA (2021-69012-35978) and New Harvest for support of this work.

## Data & resource availability

The data supporting this study are available within the article’s Supplementary files. Extra data are available from the corresponding author upon request. Immortalized cells are available from the lab for distribution for research groups, and ongoing efforts are being made to deposit the cells in a cell bank for general use.

