## Supplementary material for "Immortalized bovine satellite cells for cultured meat applications": Suppelementary Materials

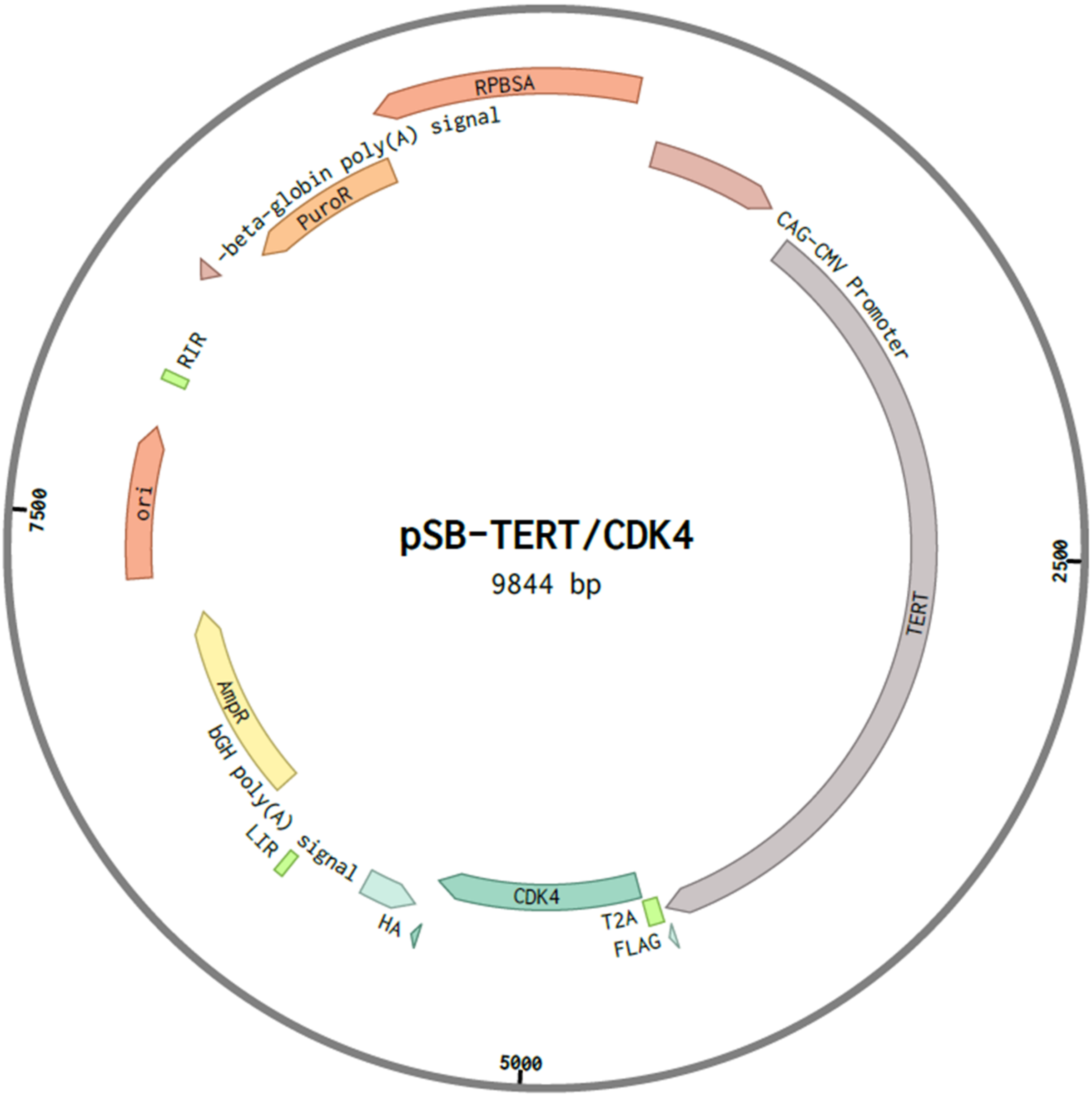


Figure S1: Sleeping beauty transposon plasmid containing TERT and CDK4 under the constitutive promotion of CMV, and a puromycin resistance under the constitutive promotion of RPBSA.

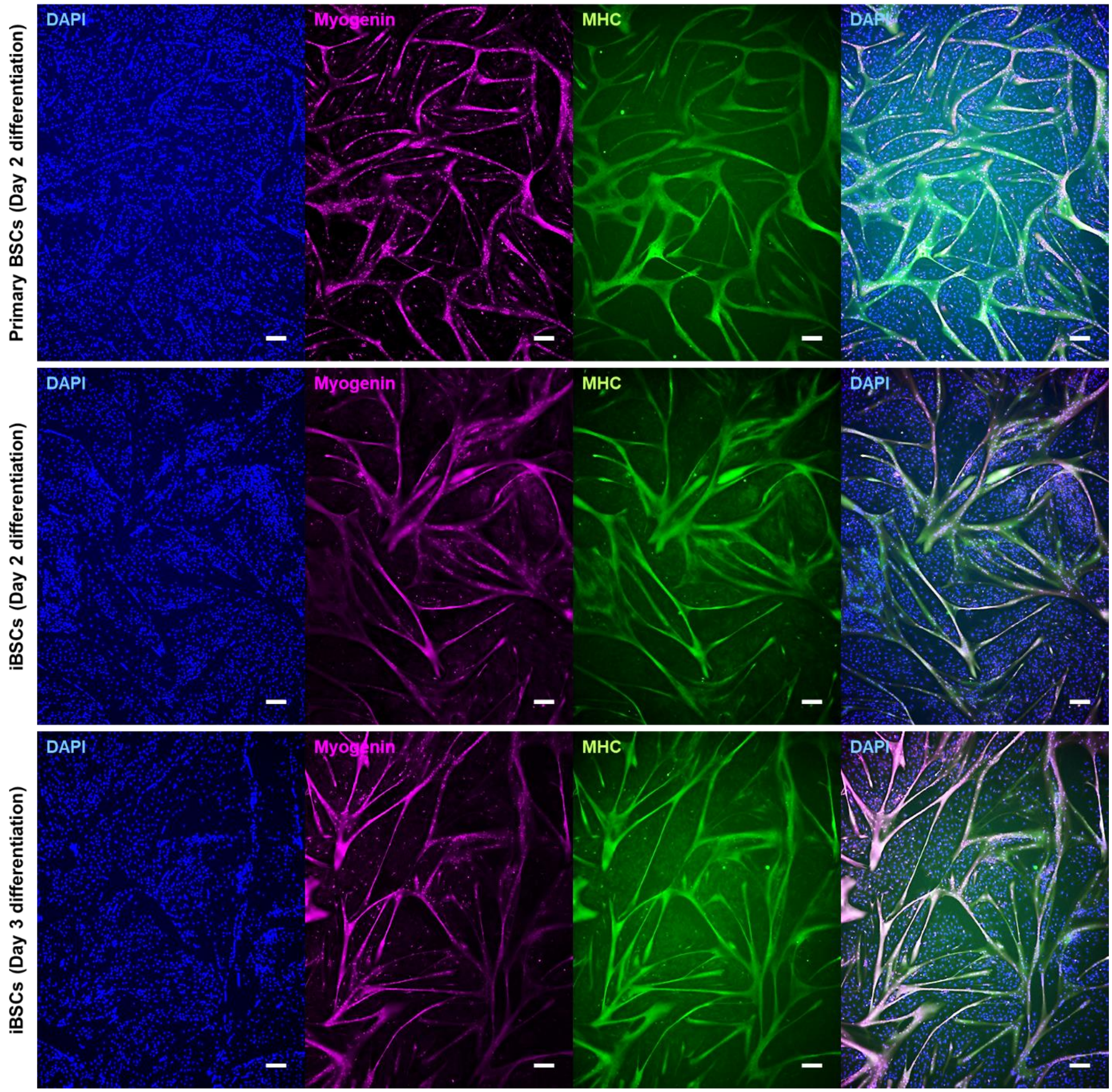


Figure S2: Immunostaining of differentiated BSCs and iBSCs. Cells were cultured to confluency and differentiated in medium containing 2% FBS for two or three days, and stained for nuclei (DAPI; blue), the early differentiation marker Myogenin (magenta), and the terminal fusion marker myosin heavy chain (MHC; green). The results showed that both BSCs and iBSCs could fuse into robust multinucleated myotubes. No clear differences in differentiated cell morphology were observed. Scale bars are 200 μm.

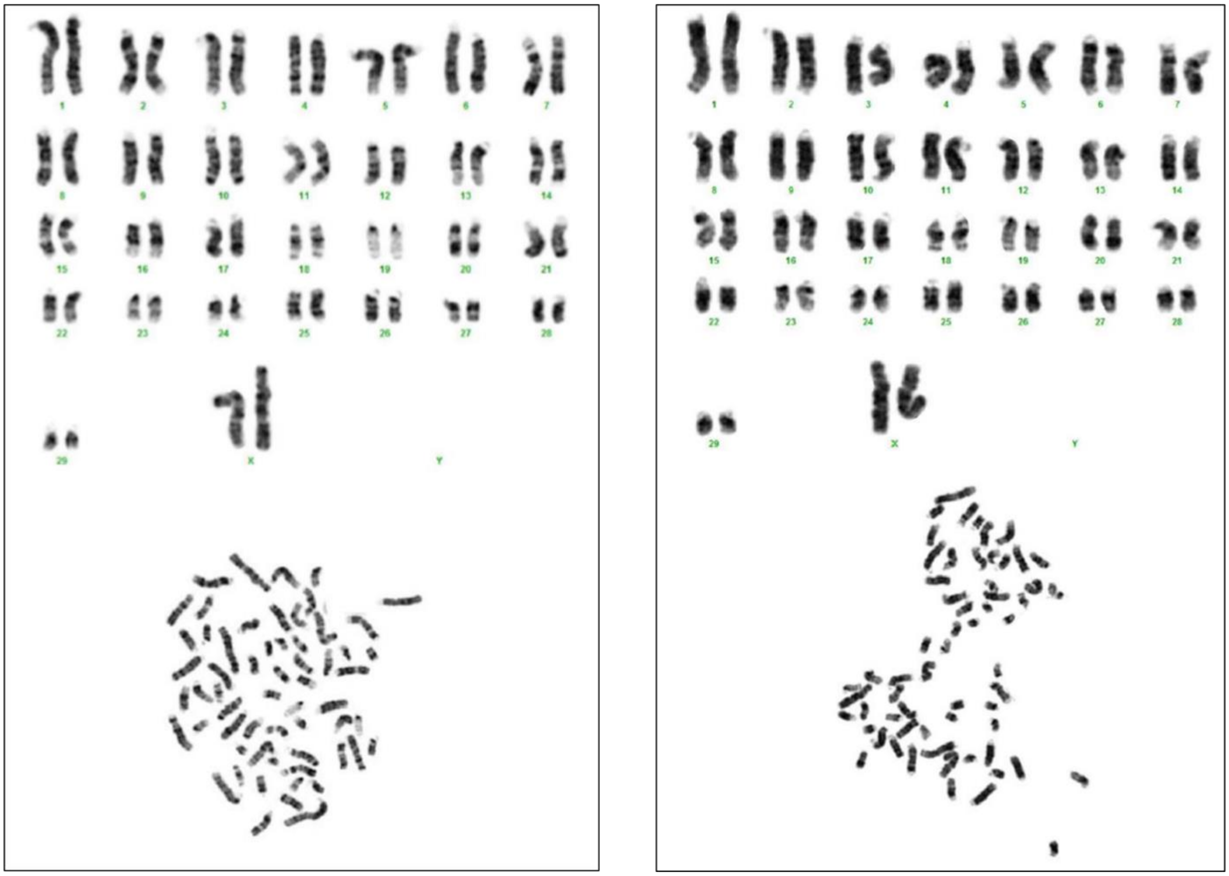

Figure S3: Karyotyping of iBSCs. Duplicate G-banded metaphase spreads of iBSCs (passage 28) revealed a normal female bovine karyotype. Lower images display metaphase chromosome spreads, while upper images display the corresponding karyotypes.
